# Radiosafe micro-computed tomography for longitudinal evaluation of murine disease models

**DOI:** 10.1101/847434

**Authors:** Nathalie Berghen, Kaat Dekoster, Eyra Marien, Jérémie Dabin, Amy Hillen, Jens Wouters, Jasmine Deferme, Thibault Vosselman, Eline Tiest, Marleen Lox, Jeroen Vanoirbeek, Ellen De Langhe, Ria Bogaerts, Marc Hoylaerts, Rik Lories, Greetje Vande Velde

## Abstract

Implementation of *in vivo* high-resolution micro-computed tomography (μCT), a powerful tool for longitudinal analysis of murine lung disease models, is hampered by the lack of data on cumulative low-dose radiation effects on the investigated disease models. We aimed to measure radiation doses and effects of repeated μCT scans, to establish cumulative radiation levels and scan protocols without relevant toxicity.

Lung metastasis, inflammation and fibrosis models and healthy mice were weekly scanned over one-month with μCT using high-resolution respiratory-gated 4D and expiration-weighted 3D protocols, comparing 5-times weekly scanned animals with controls. Radiation dose was measured by ionization chamber, optical fiberradioluminescence probe and thermoluminescent detectors in a mouse phantom. Dose effects were evaluated by *in vivo* μCT and bioluminescence imaging read-outs, gold standard endpoint evaluation and blood cell counts.

Weekly exposure to 4D μCT, dose of 540-699 mGy/scan, did not alter lung metastatic load nor affected healthy mice. We found a disease-independent decrease in circulating blood platelets and lymphocytes after repeated 4D μCT. This effect was eliminated by optimizing a 3D protocol, reducing dose to 180-233 mGy/scan while maintaining equally high-quality images.

We established μCT safety limits and protocols for weekly repeated whole-body acquisitions with proven safety for the overall health status, lung, disease process and host responses under investigation, including the radiosensitive blood cell compartment.

## Introduction

*In vivo* micro-computed tomography (μCT), an excellent technique to longitudinally evaluate disease progression in small animal models, allows non-invasive visualization of different pathogenic lung processes, (e.g. emphysema, fibrosis, lung infection and metastasis) ^1–8^. To study disease progression and therapeutic effects in real-time in individual animals, consecutive scanning is essential and enables a manifold reduction in experimental animals, which is of both ethical and economical importance. Yet, the biological effects of ionizing radiation from repeated μCT scanning remain a concern, as these are largely unexplored. Adverse radiation effects have been extensively studied for the field of radiotherapy. However, these doses (ranging from 4 to 20 Gy) and dose-rates are an order of magnitude higher than doses delivered with repetitive μCT of animal models (less than 800 mGy for a single μCT-acquisition) ^9–13^. In healthy animals, weekly or biweekly repeated respiratory-gated μCT scans for 5 to 12 weeks were well tolerated by the animals, had no radiotoxic effects on the lungs and no interference with μCT lung read-outs ^10,14^. However, radiosafety was never investigated in disease models involving rapidly dividing cells that may be differently sensitive to x-ray exposure of a μCT scan.

To exploit high-resolution μCT to its full potential and implement it in the preclinical respiratory research workflow, it is essential to investigate potential effects of repeated low-dose radiation on disease processes and host response. This is particularly relevant for radiosensitive organs such as the lungs and/or when the disease process involves rapidly dividing cells, as in many models of cancer, metastasis, inflammation and tissue remodelling ^9,15^. Currently, preclinical μCT applications focused mainly on acquiring high-resolution and - quality images while little attention was given to the delivered doses and their potential radiotoxic effects. Standard operating procedures to measure and evaluate dose and dose-effects in a preclinical setting and systematic measurements better characterizing the biological radiation effects are urgently needed. Therefore, this study assessed the effects of low-dose radiation of longitudinal whole-body μCT protocols on metastasis inflammation and fibrosis as well as host responses in murine models of lung disease. We aimed to establish high-quality generic μCT protocols that ensure safe repeated high-resolution micro-CT evaluation not interfering with animal health, the radiosensitive blood cell compartment and host response or the disease processes under investigation.

## Results

### Repetitive 4D μCT in a lung metastasis model

We investigated the potential effects of whole-body x-ray exposure from repeated μCT on lung metastasis disease in a syngeneic model of rapidly dividing cells. One group with induced lung metastasis was scanned at baseline and weekly for 4 weeks, the second group was scanned at baseline and endpoint only (Fig. 1). 4D μCT-scanning with retrospective respiratory-gating allowed acquisition of high-quality images and functional data corresponding with inspiration and expiration. No differences due to repeated μCT were found in body weight, tumour load measured by BLI, nor in μCT read-outs for metastatic burden and host response (Fig. 2 A-C). Next, we assessed the effects on the radiosensitive blood cell compartment, analysing the circulating blood cells (supplementary table S1, Fig. 2D). Weekly scanned mice showed a decreased platelet count (mean −251.2 * 10^3^ cells/μL; 95% CI: −327.4 to −175.0) and absolute white blood cell number (mean −0.4262 * 10^3^ cells/μL; 95% CI: −0.7740 to −0.0783), attributed to a decrease in absolute number of circulating lymphocytes (mean −0.2892 * 10^3^ cells/μL; 95% CI: −0.4435 to −0.1349). Furthermore, eosinophils were decreased (mean −0.0125 * 10^3^ cells/μL; 95% CI: −0.0185 to −0.0065), whereas number of neutrophils and red blood cells remained unaffected. In conclusion, repetitive 4D μCT-scanning influenced circulating blood cells in a lung metastasis model without clinical effects or change in disease outcomes.

**Figure 1:**
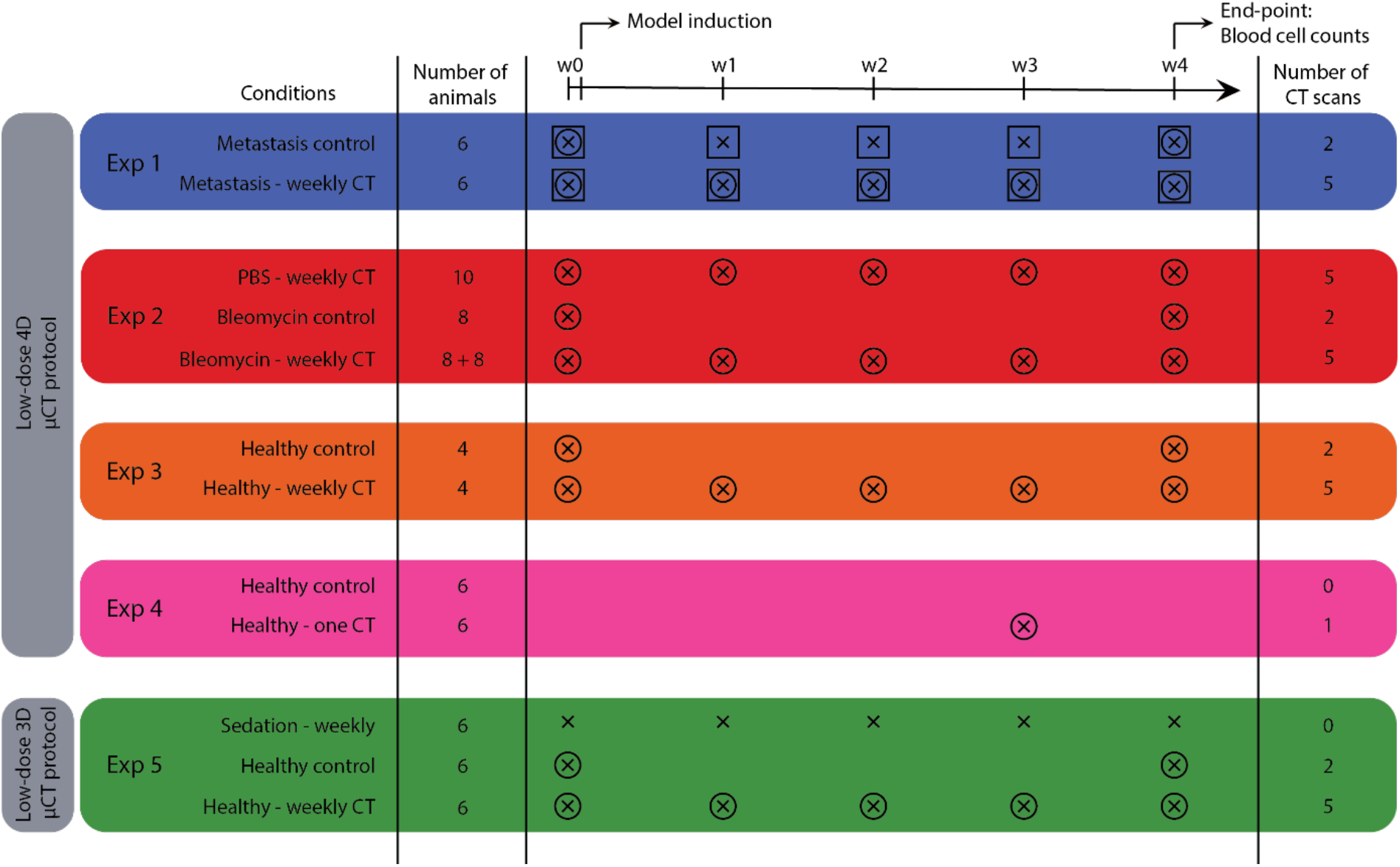
Experimental set-up. This scheme summarizes isoflurane sedation, number and timing of BLI and μCT scans, model induction and number of animals in each experimental group. **Experiment 1** compares mice with lung metastasis (DBA/2 strain) that underwent μCT-scans at baseline and weekly after metastasis induction for 4 weeks, with a metastasis group that was scanned with μCT only at baseline and endpoint. The lung metastasis burden in both groups was monitored with weekly BLI scans. **Experiment 2** compares bleomycin- (8 mice with 0.04U and 8 mice with 0.05U bleomycin) and sham-instilled mice (5 + 5 mice) of the C57Bl/6 strain that were weekly μCT-scanned with a bleomycin control group (8 mice) that was only scanned at baseline and endpoint. **Experiment 3** focuses on the effect of weekly μCT scans without the presence of disease. Analogous to experiment 1, healthy DBA/2 mice received either only a μCT scan at the beginning and at the end of the experiment or weekly scans for the entire experiment duration. In **experiment 4**, mice (C57Bl/6 strain) scanned once are compared for delayed effects after one week with mice receiving zero scans. **Experiment 5** investigates the potential effect of the low-dose 3D μCT protocol after 5 weekly μCT scans compared to one scan at the beginning and one at the end. A third control group was included that was sedated with isoflurane and handled as all other mice but did not receive any μCT scans. (x = isoflurane, ○ = μCT scan, ◻ = BLI scan)

**Figure 2:**
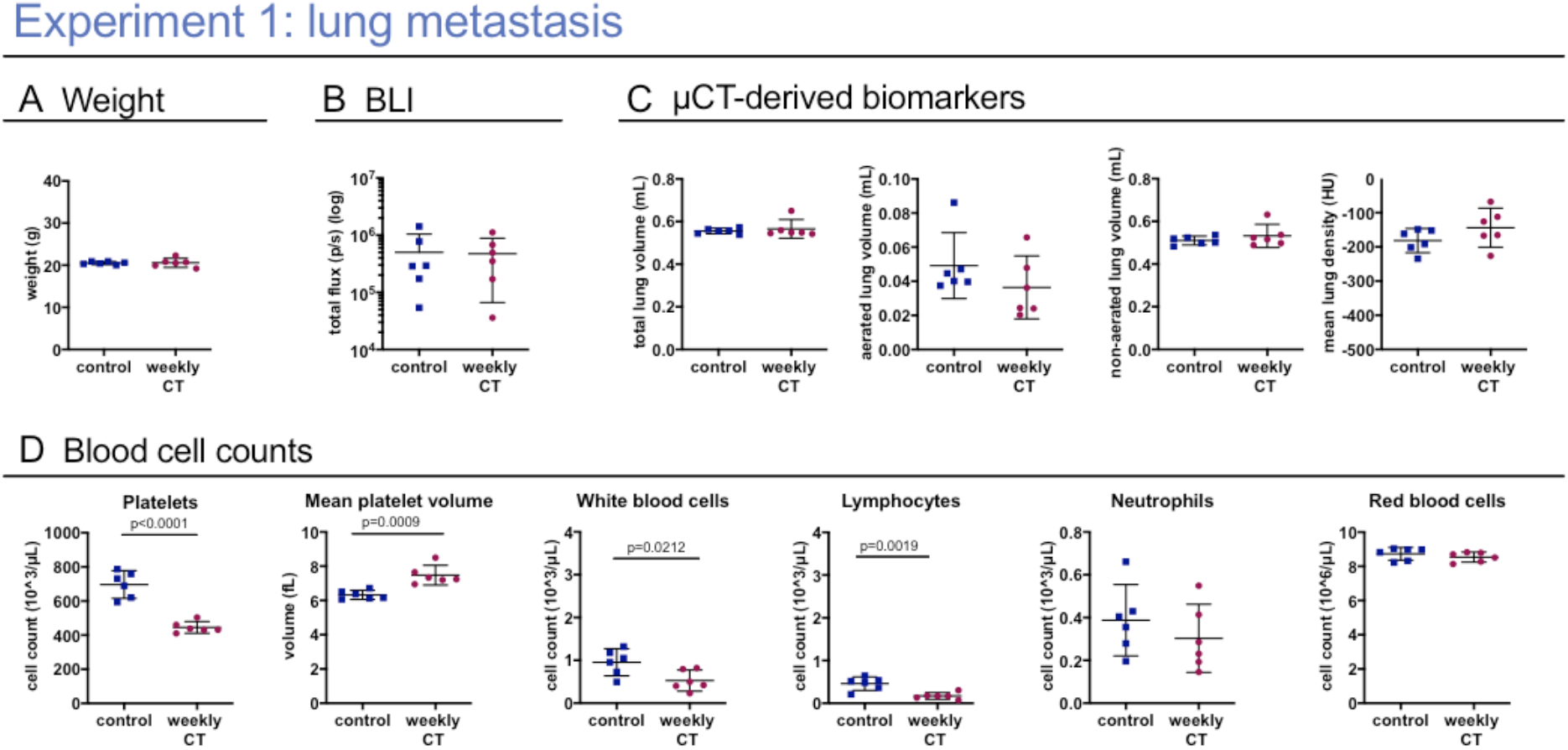
Weekly low-dose 4D μCT does not influence the general health and disease outcomes but induces a sub-clinical decrease in white blood cell and platelet counts in a murine metastasis model. **Experiment 1:** weekly-low dose 4D μCT scanning of metastasis-bearing mice induces a decrease in circulating platelets, increase in mean platelet volume, decrease in red blood cells and absolute white blood cell count, due to a decrease in number of lymphocytes. **(A)** Mouse body weight at end point. **(B)** *In vivo* BLI signal intensity expressed as total flux (p/s) from the lung, measuring metastatic load. **(C)** μCT-derived biomarkers (total lung volume, aerated lung volume, non-aerated lung volume and mean lung density. **(D)** Selected blood cell parameters: absolute platelet cell count, mean platelet volume, white blood cell count, lymphocyte count and neutrophil count and red blood cell count. Data are presented as individual values, group mean and 95% confidence intervals. P-values are presented in the graph when p < 0.05. HU, Hounsfield units.

### Repetitive 4D μCT in a lung inflammation and fibrosis model

Next, repeated low-dose radiation was evaluated in a model involving endogenous rapidly dividing cells: bleomycin-induced lung fibrosis. Mice instilled with PBS or bleomycin were scanned at baseline and weekly for 4 weeks, compared with bleomycin-instilled controls only scanned at baseline and endpoint (Fig. 1).

Due to high mortality in the bleomycin groups (2 mice of 8 (weekly CT) and 3 of 8 (control) reached the endpoint), our analysis was underpowered. Therefore, we included data of an experiment with the same set-up, except that mice were instilled with a reduced dose of bleomycin (0.04 U instead of 0.05 U). All mice were weekly scanned (Fig. 3 in grey). 3 out of the 8 mice instilled with bleomycin reached the endpoint in this second cohort.

**Figure 3:**
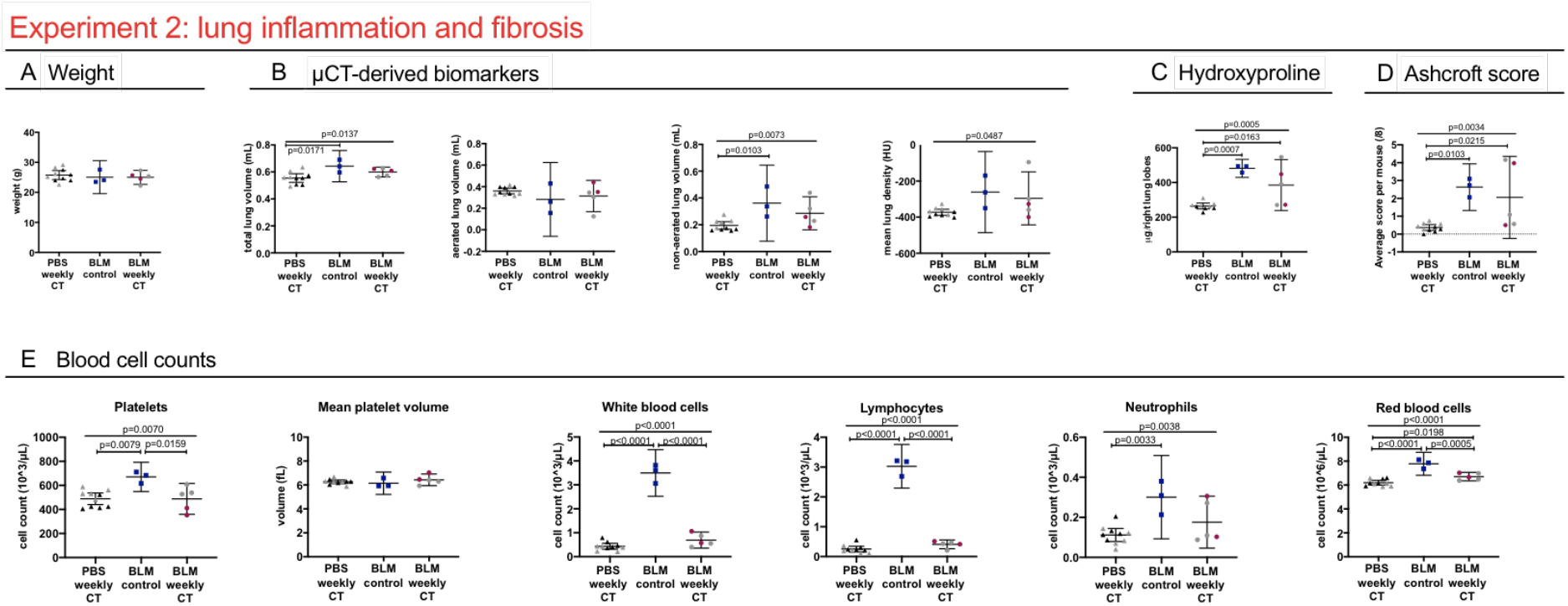
Weekly low-dose 4D μCT alters blood cell counts in a bleomycin-induced mouse model. **Experiment 2:** weekly-repeated 4D μCT scanning of bleomycin induced mice results in a decrease in platelets, red blood cells and a decrease in white blood cells, attributed to decreased lymphocyte counts. **(A)** Mouse body weight at end point **(B)** μCT-derived biomarkers reflecting disease progression of pulmonary fibrosis at endpoint (total lung volume, aerated lung volume, non-aerated lung volume and mean lung signal density). **(C)** collagen content as measured by hydroxyproline quantification of the right lung lobes and **(D)** Ashcroft score of extent of fibrosis of the left lung lobes **(E)** selected blood cell counts: absolute platelet cell count, mean platelet volume, white blood cell count, lymphocyte count and neutrophil count and red blood cell count. Data presented as individual values, group mean and 95% confidence intervals. Grey points represent mice instilled with 0.04 U of bleomycin, other points with 0.05 U. P-values and p-adjusted values are presented in the graph when p < 0.05. HU, Hounsfield unit.

Similar to the lung metastasis model, platelet counts were decreased (mean −182.7 * 10^3^ cells/μL; 95% CI: −333.8 to −31.62) in the weekly scanned bleomycin group compared to control bleomycin group (Fig. 3E, supplementary table S2). Absolute white blood cell number was lower (mean −2.805 * 10^3^ cells/μL; 95% CI: −3.277 to −2.334), with less circulating lymphocytes (mean −2.619 * 10^3^ cells/μL; 95% CI: −2.936 to −2.301). Red blood cell numbers were also decreased (mean −1.072 * 10^6^ cells/μL; 95% CI: −1.658 to −0.4862). Platelet counts for PBS-instilled weekly scanned mice were similarly decreased (mean −181.9 * 10^3^ cells/μL; 95% CI: −318.1 to −45.72) and absolute white blood cells (mean −3.064 * 10^3^ cells/μL 95% CI: −3.489 to −2.639) (attributed to a decrease in lymphocytes (mean −2.776 * 10^3^ cells/μL; 95% CI: −3.062 to −2.490)) and less red blood cells (mean −1.586 * 10^6^ cells/μL 95% CI: −2.114 to −1.058).

Although the analysis of differences in disease severity between the bleomycin control and weekly scanned mice remained underpowered, no changes were found between these groups concerning body weight and our data indicate no clear influence from repeated scanning on mortality or pathology (extent of lung inflammation and fibrosis, assessed by histology, collagen content and μCT read-outs (Fig. 3A-D)). PBS-instilled control mice were unaffected by radiation as evaluated by histology and μCT.

In conclusion, repetitive 4D μCT-scanning lowered circulating blood cells, irrespective of disease status.

### Repeated 4D μCT in healthy mice

To further elucidate the contribution of disease status to the effects of repeated μCT, we investigated healthy mice. One group was scanned with the weekly regime (n = 4) and the other only at beginning and endpoint (n = 4). No differences were found concerning general health and lung μCT read-outs (Fig. 4A). Similar to the findings in the two disease models, a decrease in platelets (mean −387.9 * 10^3^ cells/μL; 95% CI: −521.6 to −254.3) and white blood cells (mean −0.3510 * 10^3^ cells/μL; 95% CI: −0.6859 to −0.01608) (attributed to a decrease in lymphocytes (mean −0.02849 * 10^3^ cells/μL; 95% CI: −0.5149 to −0.05496)) and a decrease in red blood cells (mean −1.832 * 10^6^ cells/μL; 95% CI: −3.637 to −0.02679) was found in the weekly scanned healthy mice compared to controls (Fig. 4B, supplementary table S3). These results confirm the effect of repeated low-dose 4D μCT-scanning on circulating blood cells, irrespective of disease or inflammatory status.

**Figure 4:**
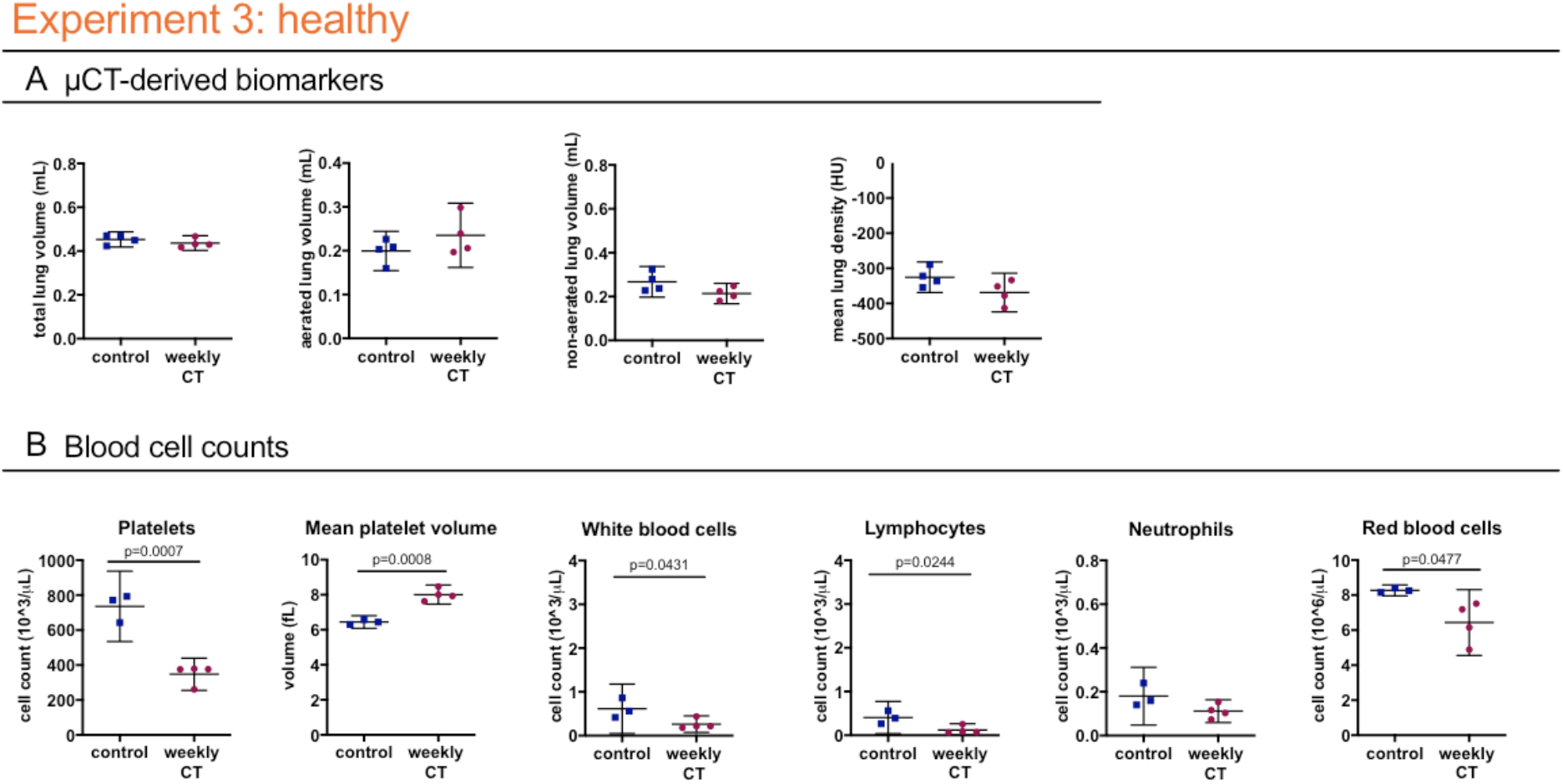
Weekly low-dose 4D μCT does not influence the general health and disease outcomes but induces a decrease in blood cell counts of healthy mice. **Experiment 3:** weekly-low dose 4D μCT scanning of healthy mice induces a decrease in platelets, increase in mean platelet volume, decrease in red blood cells and white blood cells, attributed to decreased lymphocyte counts. **(A)** μCT-derived biomarkers show no differences in healthy mice at endpoint (total lung volume, aerated lung volume, non-aerated lung volume and mean lung density). **(B)** Blood cell counts: absolute platelet cell count, mean platelet volume, white blood cell count, lymphocyte count and neutrophil count and red blood cell count. Data presented as individual values, group median and 95% confidence intervals. P-values are presented in the graph when p < 0.05. HU, Hounsfield unit.

To exclude that lower blood cell counts were a delayed effect of the second last scan, we compared blood cell counts of mice receiving a single scan a week before sacrifice (n = 6), and mice receiving no scan at all (n = 6). No significant differences were found for platelets, white blood cells and lymphocytes (Fig. 5, supplementary table S4), indicating the observed effects in the previous experiments were indeed related to repeated exposure.

**Figure 5:**
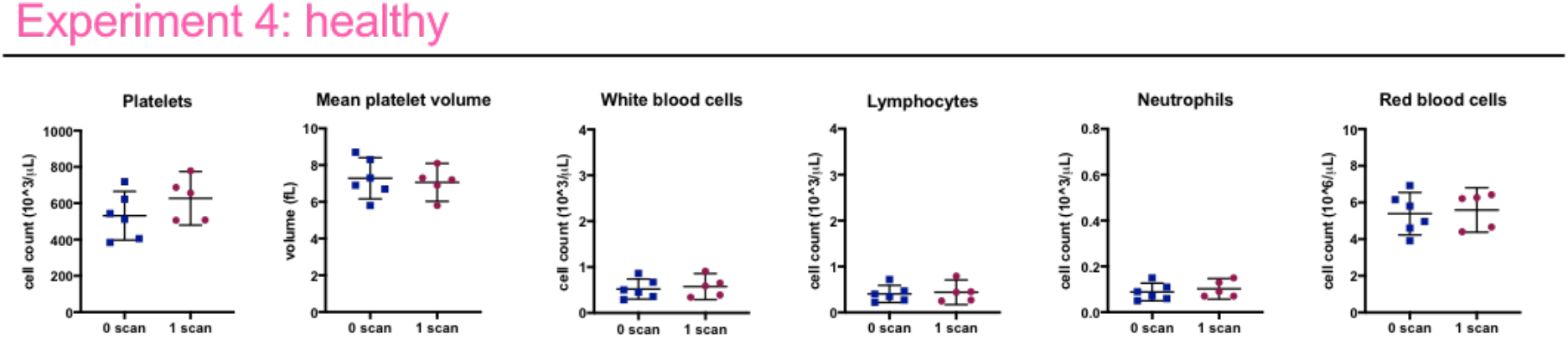
a low-dose 4D μCT scan does not affect blood cell counts one week after scanning. **Experiment 4:** No differences are found in circulating blood cell counts between healthy control and scanned mice at endpoint, i.e. 1 week after the scan. Data presented as individual values, group mean and 95% confidence intervals. P-values are presented in the graph when p<0.05.

To isolate effects of repetitive x-ray exposure from potential influence of stress and anaesthesia associated with μCT, we included an additional healthy control group subjected to weekly anaesthesia and handling without μCT. This group showed no differences in blood cell counts (Fig. 6B-D), further confirming that blood cell count effects can be attributed to repeated x-ray exposure.

**Figure 6:**
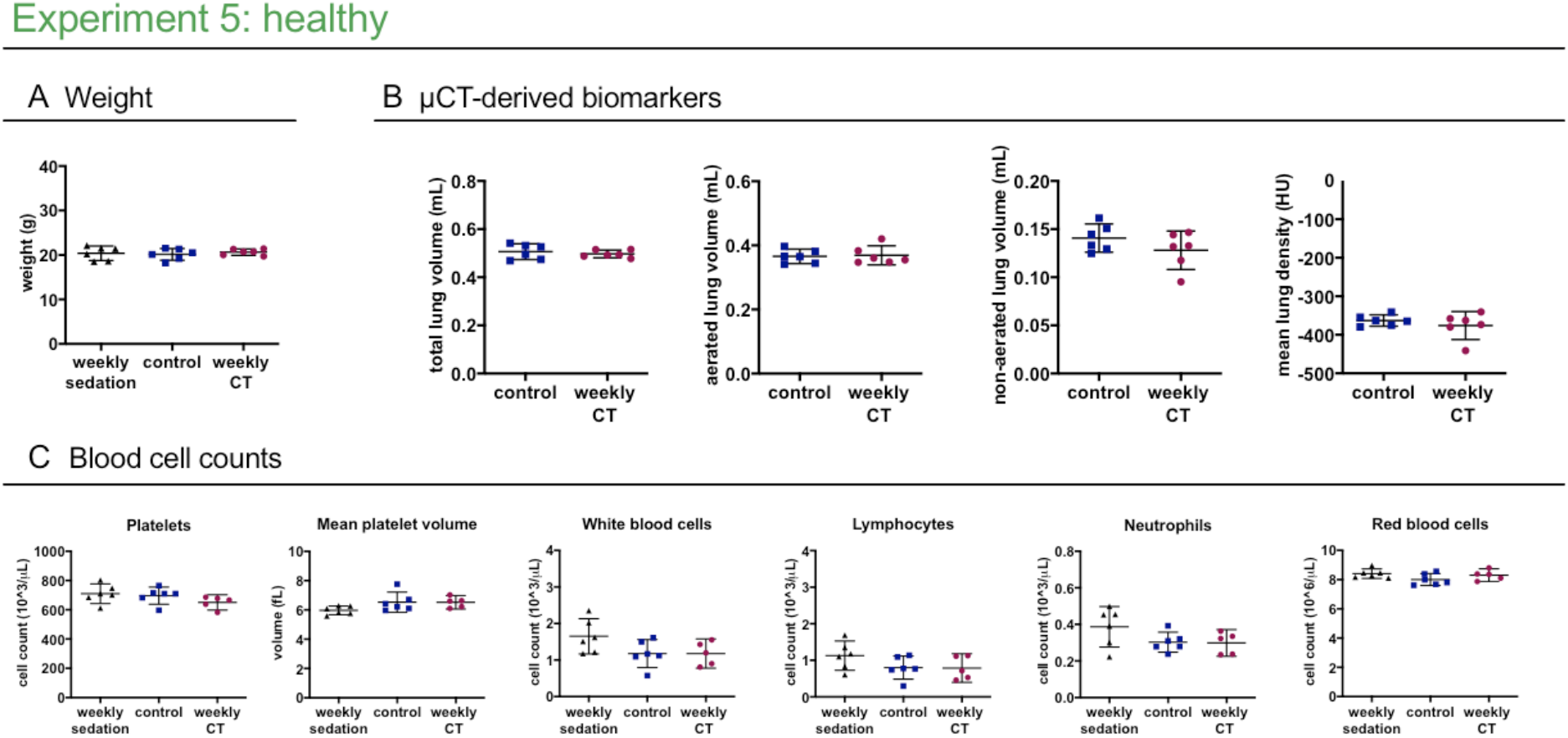
Weekly low-dose 3D μCT is devoid of any effects on health, lung readouts and circulating blood cell counts. **Experiment 5** compares healthy mice scanned weekly or only at baseline and endpoint, and mice that underwent weekly isoflurane anaesthetics and handling without undergoing any μCT scans to isolate a potential effect from stress and anaesthesia from an effect of the x-ray dose associated with a μCT scan (weekly sedation). **(A)** Mouse body weight at end point. **(B)** μCT-derived biomarkers show no difference at endpoint (total lung volume, aerated lung volume, non-aerated lung volume and mean lung density) between the healthy control and healthy weekly scanned group. **(C)** selected blood cell counts: weekly low-dose 3D μCT scanning or weekly isoflurane sedation does not change the platelet, white blood cell or red blood cell counts. Data presented as individual values, group mean and 95% confidence intervals. P-values and p-adjusted values are presented in the graph when p<0.05.

### Repeated 3D μCT: image quality and circulating blood cells

To eliminate effects on blood cells, while retaining high-quality images, we optimized a 3D protocol with expiration-weighted gating but without the possibility to acquire functional data. Scan time and dose were hereby much reduced compared to retrospective respiratory-gated 4D μCT. For accurate dosimetry, we used three different methods (table 1). Radiation dose of a single μCT scan with the 3D versus 4D protocol ranged between 180-233 mGy versus 540-699 mGy respectively.

Our 3D expiration-weighted strategy did not introduce any movement artefacts compared to respiratory-gated 4D scans. Analysis of signal-to-noise (SNR), contrast-to-noise ratio (CNR) and visual inspection confirmed equally high image quality (Fig. 7) (a mean SNR of 26.9 and 25.65 and a mean CNR of 26.13 and 24.76 for 4D and 3D protocols, respectively, defined by the Rose criterion ^16,17^.

**Table 1:** dose measurements for the low-dose 4D and 3D μCT protocols. Dose measurements were performed with ionization chamber (IC) in air, thermoluminescent detectors (TLDs) in mouse phantom and optical fiber radioluminescence (RL) probe in mouse phantom as specified in materials and methods.

| | Low-dose 4D<br>μCT protocol | Low-dose 3D<br>μCT protocol |
| --- | --- | --- |
| IC in air | 540 mGy | 180 mGy |
| TLDs in phantom | 699 mGy | 233 mGy |
| RL probe in phantom | 585 mGy | 195 mGy |

**Figure 7:**
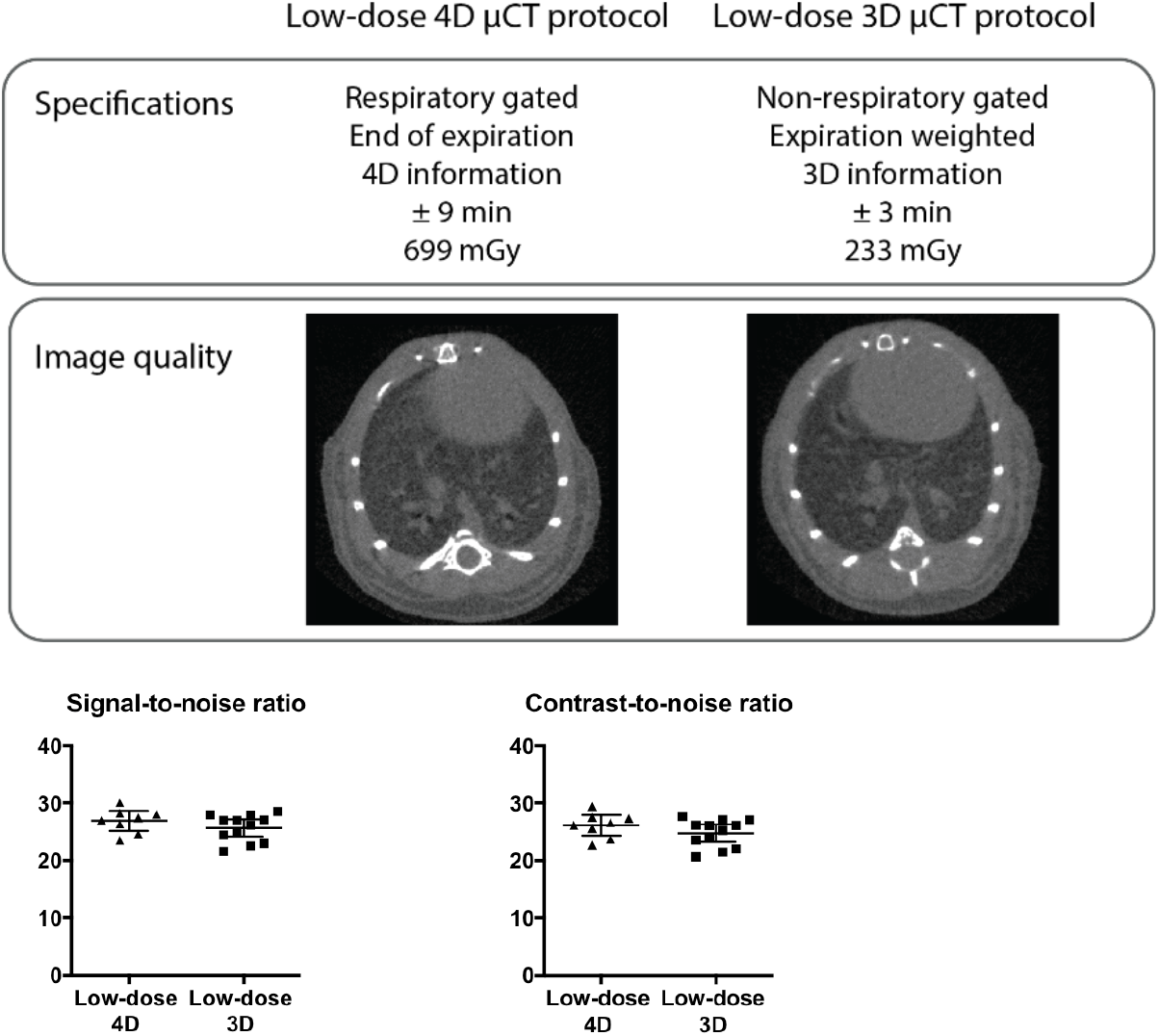
Low-dose expiration-weighted 3D μCT yields equally high image quality as a 4D respiratory-gated protocol. **(A)** Specifications of the 3D and 4D imaging protocols compared with representative reconstructed tomographic images at the level of lung and heart for both protocols, along with graphs of **(B)** signal-to-noise ratio and contrast-to-noise ratio compared for the respiratory-gated 4D (n = 8) and the expiration-weighted 3D μCT protocol (n = 12). Data presented as individual values, group mean and 95% confidence intervals.

In healthy mice, weekly 3D μCT did not affect body weight, μCT-derived lung biomarkers nor blood cell counts (Fig. 6, supplementary table S5): platelet numbers, white blood cells (lymphocytes as well as neutrophils) and red blood cells in weekly scanned healthy mice were not affected. In conclusion, repeated low-dose 3D μCT has no radiation effects on lungs or blood cells, while maintaining high image quality even for lung imaging in free-breathing mice.

## Discussion

We investigated radiation dose and effects of repetitive whole-body μCT in different murine lung disease models and healthy mice, using a longitudinal imaging set-up typical for preclinical *in vivo* experiments: baseline scan, 4 weekly μCT scans for disease progression and sacrifice after last scan for *ex vivo* read-outs. Using a low-dose 4D retrospective respiratory-gating protocol, we observed no effects of repetitive scanning on disease outcomes in mice with lung metastasis or healthy mice, but detected a decrease in circulating blood platelets and lymphocytes. With a 3D expiration-weighted scanning protocol, reducing the radiation dose by two thirds, this effect could be eliminated whilst retaining equally high imaging quality.

We analysed both healthy and diseased DBA/2 and C57Bl/6 mice, commonly utilized strains in preclinical pulmonary research. As we used immunocompetent mice, we can conclude that μCT has the potential to study disease processes where host response is an important factor, with C57Bl/6 mice in particular susceptible to lung fibrosis ^18^. As we did not observe nor quantify an effect of radiation after weekly 4D μCT in healthy animals, our results are in line with previous studies reporting the absence of radiation-induced lung or cardiac damage in healthy mice scanned weekly for 5 to 12 weeks ^10,14^. Furthermore, we assessed different lung disease models involving rapidly dividing exogenous and endogenous cells in immunocompetent mice. Repetitive μCT had no influence on disease progression in mice with progressing lung metastases. In mice with lung inflammation and fibrosis, our data also indicate no clear influence from repeated scanning on fibrosis, although this experiment was underpowered. Repeated low-dose 4D μCT can therefore be considered safe for the animal and disease process under investigation, even when involving rapidly dividing cells, but we continue to recommend the use of a similarly radiation exposed control group.

We next investigated whether repeated low-dose x-ray exposure may still have subtle effects on more radiosensitive processes by analysing circulating blood cells since the hematopoietic system and lymphocytes in particular are very sensitive to radiotoxicity ^19,20^. In repetitively scanned mice, we observed less (but larger) circulating platelet and lymphocytes, neutrophils remained unaffected. Red blood cells numbers were lowered in repetitively scanned healthy and bleomycin-induced C57BL/6 mice, but not in DBA/2 mice with or without metastasis. The lower platelet and lymphocyte counts after repeated radiation in mice with lung metastasis, fibrosis and in healthy mice, points to disease-independent effect of the cumulative radiation dose. The decrease in platelets is likely sub-clinical, since we detected no bleeding problems. Also a murine platelet reduction up to 70% is reported not to result in clinical effects ^21^. Defining values of clinically significant lymphopenia is more difficult, given the high variability in reported murine reference values ^22^. The lymphocyte reduction could be sub-clinical, not affecting the immune system, given the absence of infections or a correlation with disease outcomes in our study. Yet, we cannot formally exclude minimal effects on immune system functionality. Importantly, we nevertheless saw no influence of the decreased platelets and lymphocytes on the studied disease processes.

We examined blood cell counts immediately after the last scan (typical set-up of last scan followed by sacrifice and *ex vivo* work-up). Therefore, results reflect the short-term effect of cumulated radiation from weekly repeated μCT scans, no conclusions can be drawn on long-term effects and potential recuperation with time. Noteworthy, blood cell changes were not an effect of the last scan, since control groups received the same endpoint scan. No changes were seen one week after a single scan, ruling out delayed effects of the second-last scan. Moreover, no differences in circulating blood cells were observed comparing μCT-scanned healthy mice to controls that were sedated but never scanned. Therefore, we conclude that effects result from the cumulative radiation exposure.

To design longitudinal μCT protocols without radiotoxicity and still considering breathing movement corrections, we developed a respiration-weighted 3D protocol. This could lower the radiation dose per scan with two-thirds while maintaining the same lung image quality as the 4D protocol, with no increased movement artefacts, but without the possibility to extract functional lung read-outs (e.g. tidal volume, readouts at inspiration). This 3D protocol is nevertheless useful for most studies, where volume data from inspiratory phase are not needed. With this 3D protocol, we could eliminate all previously found radiation effects on blood cell counts, thereby offering a generically applicable longitudinal μCT protocol with demonstrated safety for the animal and (patho-)physiological processes under investigation. Our results further expand knowledge about maximum acceptable repeated dose exposure, as previous reports found no blood cell count changes in C57Bl/6 mice scanned less frequently (every other week for 3 times) ^23^, as well as in mice scanned more frequently (3 times/week for 4 weeks) ^24^, but with markedly lower radiation doses than our 3D protocol (reported dose 16.19 mGy measured at phantom center versus 180-233 mGy dose).

Indispensable in guarding over radiation exposure is awareness of dose exposure and hence accurate dosimetry. As currently no standard operating procedures exist for preclinical μCT dosimetry, we measured radiation dose of our 4D and 3D μCT protocols by three different methods: IC in air, in-phantom TLDs and an in-phantom RL probe. The doses measured in-phantom were higher compared to in-air, as expected due to the contribution of scattered x-rays. The IC is the reference and used for calibration of TLDs and RL probe. Nevertheless, size limits its use in μCT, since its dimensions do not enable complete dose profile measurements and the closed lead shielding may hamper insertion of an IC in the field-of-view during scanning. Moreover, the interest of using such IC combined with a mouse phantom is limited in contrast with conventional CT in the clinical context where the dimensions of the IC are negligible compared to those of the CT Dose Index phantoms. TLDs are a practical alternative. Although their energy response is not as good as the IC, they are made of tissue-equivalent material and small enough to be inserted in-phantom. An inherent limitation of TLDs is they do not allow real-time dosimetry, needed during preclinical *in vivo* scan parameter optimization. Compared to TLDs, the RL probe results are subject to higher uncertainty because of the higher energy dependence of the RL material. Nevertheless, the advantages for μCT are its small size and capacity to perform online and real-time measurements, useful for scan protocol optimization.

To summarize, it is necessary and within our potential to design high-quality and safe μCT protocols, regarding the murine health status, disease process and host responses under investigation. We have established an upper dose limit to be delivered with repeated μCT scanning: a dose of 540-699 mGy delivered weekly for 5 times, can be considered as physiologically safe with a sub-clinical drop in circulating blood cell counts, while a dose of 180-233 mGy per single scan delivered under the same longitudinal regime is safe in absolute terms. More specifically, our results indicate the possibility to design high-resolution μCT protocols without influence on the most radiosensitive processes in the body, thereby ideal to study (lung) disease processes and host responses in rodent models.

## Materials and methods

### Animals

Mice were kept in individually ventilated cages or filter top cages with free access to food and water in a conventional animal facility. The syngeneic mouse model of lung metastasis was induced by tail vein injection of cells from the squamous cell carcinoma (SCC) lung cancer cell line KLN205 (10^5^ cells in 200 μl PBS) in 8-week-old female DBA/2 mice under transient isoflurane (2% in oxygen) gas anaesthesia (Envigo, Venray, The Netherlands) ^1^. For the bleomycin-induced lung inflammation and fibrosis model, 8-week-old male C57Bl/6 mice (Janvier, Le Genest, France) were anesthetised with a mixture of ketamin (Nimatek 10 mg/ml, Europet, Gemert-Bakel, The Netherlands) and xylazine (2% Xyl-M 1 mg/ml, VMD, Arendonk, Belgium). Via a tracheotomy, 50 μL of Bleomycin (0.04 U or 0.05 U, Sanofi-Aventis, Diegem, Belgium) or vehicle phosphate buffered saline (PBS, Lonza, Basel, Switzerland) for sham controls was instilled ^2,3^. Mouse body weights were recorded at baseline and at least once weekly until sacrifice. For experiments conducted with healthy animals, 8-week-old female DBA/2 or male C57Bl/6 mice (Janvier, Le Genest, France) were used. An overview of the number of animals per experimental group can be found in the experimental set-up (Fig. 1). European, national and institutional guidelines for animal welfare and experimental conduct were followed (The KU Leuven Ethical Committee for animal research approved all experiments: p039/2014, p037/2017 and p227/2013).

### Bioluminescence imaging

For BLI and quantification of lung metastasis burden, an IVIS Spectrum system (CaliperLS; Perkin-Elmer, Hopkinton, MA, USA) was used with software provided by the manufacturer (Living Image version 4.4.17504). D-luciferin (in PBS, 126 mg/kg) was injected intraperitoneally, acquisition of consecutive frames was started immediately thereafter until maximum signal intensity was reached, measured as photon flux per second through a region of interest (2.9 cm × 1.8 cm) covering the lungs. Image acquisition numbers and times varied between 10 and 15 frames of 30-60s each, depending on optimal acquisition settings as a function of signal intensity intrinsic to lung metastasis grade.

### Micro-computed Tomography

Mice were anesthetized by isoflurane (1.5-2% in oxygen, Piramal Healthycare, Morpeth, Northumberlang, United Kingdom) and scanned in supine position using *in vivo* μCT (Skyscan1278, Bruker micro-CT, Kontich, Belgium) with following parameters: 50 kVp X-ray source voltage, 918 μA current, 1 mm aluminium X-ray filter, 55 ms exposure time per projection, acquiring projections with 0.9° increments over a total 220° angle, 10 cm field of view covering the whole body producing reconstructed data sets with 50 μm isotropic voxel size either with (‘4D protocol’) or without retrospective respiratory gating (‘3D protocol’). For the 4D protocol, images were acquired in list mode, with nine projections per view, logged simultaneously with the breathing cycle of the mouse and retrospectively time-based sorted, resulting in four reconstructed 3D data sets corresponding to four different breathing cycle phases (4D) (end-inspiratory, end-expiratory and two intermediate phases). 3D datasets were acquired without respiratory gating using similar settings as above, acquiring and averaging three projections per view.

Software provided by the manufacturer (TSort, NRecon, DataViewer, and CTan) was used to retrospectively gate, reconstruct, visualize, and process μCT data ^25^. For Hounsfield unit (HU) calibration, a phantom of an air-filled 1.5 mL tube inside a water-filled 50 mL tube was scanned. Based on full stack histograms of a volume-of-interest (VOI) containing only water or air, the main grayscale index of water (93) set at 0 HU and grayscale index of air (0) at - 1000 HU. Quantification of mean lung density (in HU), non-aerated lung volume, aerated lung volume, and total lung volume was carried out for a VOI covering the lung, comprising of regions of interest that were manually delineated on the coronal μCT images, thereby avoiding heart and main blood vessels. The threshold to distinguish aerated from non-aerated lung tissue volume, manually set at −287.5 HU, was kept constant for all data sets.

### Dosimetry

The radiation dose of an *in vivo* μCT scan was experimentally assessed with (1) an ionization chamber, (2) an optical fiber radioluminescence (RL) probe and (3) thermoluminescent detectors (TLDs) in a mouse phantom.

A Farmer-type ionization chamber FC65-G (IBA, Schwarzenbruck, Germany) was positioned in air at the centre of the gantry, only supported by a piece of tape placed on the top of the examination bed.

Ten MCP-N thermoluminescent detectors (TLD) (LiF:Mg, Cu, P material, Institute of Nuclear Physics, Krakow, Poland), were inserted in as many dedicated cavities in the centre of a cylindrical polymethyl methacrylate phantom (100 mm long, 20 mm diameter). The phantom was positioned at the centre of the gantry on the examination bed. Reported dose is the average over the 10 positions.

The optical fiber radioluminescence (RL) probe was inserted in the centre of a dedicated cylindrical polymethyl methacrylate phantom (100 mm long, 20 mm diameter); position of the phantom in the gantry was identical to the position of the TLD phantom.

The ionization chamber was calibrated free in air in a RQR3* reference field ^26^ at the second-standard calibration laboratory of the Belgian Nuclear Research Centre (Mol, Belgium). The TLDs and the RL probe were calibrated in air against the ionization chamber using the LD3 beam quality.

### μCT image quality

Contrast-to-noise ratio (CNR) and signal-to-noise ratio (SNR) are based on the average pixel value of the heart and calculated according to the following equations:

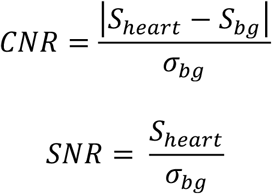

S = signal

bg = background

### Blood cell counts

Blood obtained by cardiac puncture at sacrifice, mixed with sodium citrate 3.8%, was analysed using a Cell-dyn 3700 (Abbott, Illinois, USA). Supplementary tables S1-S5 show all analysed parameters.

### Histopathology

Formalin-fixed and paraffin-embedded lung sections were stained with haematoxylin-eosin. Pulmonary fibrosis was scored using the semi-quantitative Ashcroft score ^27^. Collagen content was assessed by hydroxyproline quantification on the right lung lobes (experiment 2), as previously described ^28^.

### Statistical analysis

All measurements are reported as individual value, mean and 95% confidence intervals (CI). Data were analysed using GraphPad Prism 7.0a (Graphpad Software Inc, San Diego, CA). Based on prior work and the nature of the biological data a normal distribution was assumed. Residuals and QQ graphs were used for visual assessment of the distribution. Where of interest, groups were compared by t-test or one-way ANOVA with Bonferroni corrected multiple comparisons. Resulting differences between the means are reported with 95% confidence intervals and exact p-values.

## Acknowledgements

This research was supported by KU Leuven Internal Funds (C24/17/061 & STG/15/024). NB and KD received a PhD fellowship from the Flemish research foundation FWO (11ZP518N, 1S77319N).

## Author contribution

G.V.V. conceived the study; N.B., K.D., J.D., E.D.L., R.B., M.H., R.L. and G.V.V. designed the experiments. N.B., K.D., E.M., J.D., A.H., J.W., J.D., T.V., E.T., M.L., R.B and G.V.V. performed the experiments. N.B., K.D., J.D., T.V. and E.T. analyzed the data. M.H., E.D.L., R.L. and G.V.V. supervised the experiments and data analysis. N.B. and K.D. wrote the manuscript. J.V., R.L. and G.V.V. reviewed the manuscript. All authors have read and approved the final manuscript.

