## Supplemental tables for "Radiosafe micro-computed tomography for longitudinal evaluation of murine disease models"

**TITLE PAGE**

**Title:**

|  | **Exp1 - lung metastasis** | | | | | | |
| --- | --- | --- | --- | --- | --- | --- | --- |
|  | **Metastasis control** | | **Metastasis weekly CT** | |  | | |
|  | **mean** | **95% CI** | **mean** | **95% CI** | **p-value** | **mean difference** | **95% CI** |
| **WBC (10³/µL)** | 0.9548 | 0.6381 to 1.272 | 0.5287 | 0.2823 to 0.7751 | **0.0212** | **-0.4262** | -0.7740 to -0.0783 |
| **NEU (10³/µL)** | 0.3878 | 0.2210 to 0.5547 | 0.3032 | 0.1440 to 0.4624 | 0.3675 | -0.0847 | -0.2845 to 0.1152 |
| **NEU%** | 40.03 | 34.74 to 45.33 | 57.15 | 48.85 to 65.45 | **0.0012** | **17.12** | 8.579 to 25.65 |
| **LYM (10³/µL)** | 0.4602 | 0.3009 to 0.6194 | 0.1710 | 0.0091 to 0.2506 | **0.0019** | **-0.2892** | -0.4435 to -0.1349 |
| **LYM%** | 48.27 | 40.72 to 55.81 | 33.58 | 24.6 to 42.57 | **0.0092** | **-14.68** | -24.85 to -4.512 |
| **MONO (10³/µL)** | 0.0633 | 0.0393 to 0.0873 | 0.0355 | 0.0093 to 0.0617 | 0.0719 | -0.0278 | -0.0587 to 0.0030 |
| **MONO%** | 7,118 | 3.856 to 10.38 | 6,148 | 2.655 to 9.642 | 0.6133 | -0.97 | -5.114 to 3.174 |
| **EOS (10³/µL)** | 0.0162 | 0.0095 to 0.0228 | 0.0037 | 0.0017 to 0.0056 | **0.0009** | **-0.0125** | -0.0185 to -0.0065 |
| **EOS%** | 1,677 | 1.175 to 2.178 | 0.6867 | 0.398 to 0.9753 | **0.0013** | **-0.99** | -1.492 to -0.4883 |
| **BASO (10³/µL)** | 0.0272 | 0.0166 to 0.0377 | 0.0152 | 0.0008 to 0.0296 | 0.1146 | -0.012 | -0.0275 to 0.0003 |
| **BASO%** | 2.922 | 2.112 to 3.731 | 2,439 | 0.503 to 4.375 | 0.5676 | -0.4825 | -2.302 to 1.337 |
| **RBC (10^6^/µL)** | 8,728 | 8.351 to 9.106 | 8.54 | 8.245 to 8.835 | 0.3361 | -0.1883 | -0.6036 to 0.227 |
| **Hgb (g/dL)** | 11.78 | 11.25 to 12.31 | 11.68 | 11.32 to 12.05 | 0.6977 | -0.1 | -0.6573 to 0.4573 |
| **HCT%** | 64.6 | 61.61 to 67.59 | 64.45 | 62.37 to 66.53 | 0.9178 | -0.15 | -3.308 to 3.008 |
| **MCV (fL)** | 74 | 73.22 to 74.78 | 75.5 | 74.91 to 76.09 | **0.0028** | **1.5** | 0.6506 to 2.349 |
| **MCH (pg)** | 13.5 | 13.39 to 13.61 | 13.7 | 13.59 to 13.81 | **0.0101** | **0.2** | 0.0591 to 0.341 |
| **MCHC (g/dL)** | 18.25 | 18.19 to 18.31 | 18.13 | 18.02 to 18.24 | **0.0346** | **-0.1167** | -0.2230 to -0.0103 |
| **RDW %CV** | 16.97 | 15.8 to 18.13 | 19.78 | 18.52 to 21.04 | **0.0018** | **2,817** | 1.331 to 4.303 |
| **PLT (10³/µL)** | 696.7 | 615.8 to 777.5 | 445.5 | 411.0 to 480 | **<0.0001** | **-251.2** | -327.4 to -175.0 |
| **MPV (fL)** | 6,327 | 6.06 to 6.593 | 7,478 | 6.902 to 8.055 | **0.0009** | **1,152** | 0.6012 to 1.702 |
| **PCT %** | 0.4415 | 0.3804 to 0.5026 | 0.3328 | 0.3015 to 0.3642 | **0.0022** | **-0.1087** | -0.1681 to -0.0492 |
| **PDW 10(GSD)** | 17.6 | 16.95 to 18.25 | 19.57 | 18.79 to 20.35 | **0.0006** | **1,967** | 1.087 to 2.846 |

**Supplementary table 1 S1: blood cell counts of control and weekly scanned mice with induced lung metastasis.**

WBC=white blood cells; NEU=neutrophils; LYM=lymphocytes; MONO=monocytes; EOS=eosinophils; BASO=basophils; RBC=red blood cells; Hgb=hemoglobin; HCT=hematocrit; MCV=mean corpuscular (cell) volume; MCH=mean corpuscular hemoglobin; MCHC=mean corpuscular hemoglobin concentration; RDW=red cell distribution width; PLT=platelets; MPV=mean platelet volume; PCT=plateletcrit

|  | **Exp2 - lung inflammation and fibrosis** | | | | | | | | | | | | | | | |
| --- | --- | --- | --- | --- | --- | --- | --- | --- | --- | --- | --- | --- | --- | --- | --- | --- |
|  | **PBS weekly CT** | | **Bleomycin control** | | **Bleomycin**  **weekly CT** | |  | **PBS vs BLM CO** | | | **PBS vs BLM weekly CT** | | | **BLM CO vs BLM weekly CT** | | |
|  | **mean** | **95% CI** | **mean** | **95% CI** | **mean** | **95% CI** | **p-value** | **p-adj value** | **mean difference** | **95% CI** | **p-adj value** | **mean difference** | **95% CI** | **p-adj value** | **mean difference** | **95% CI** |
| **WBC (10³/µL)** | 0.4327 | 0.3100 to 0.5554 | 3.497 | 2.522 to 4.472 | 0.6912 | 0.3573 to 1.025 | **<0.0001** | **<0.0001** | -3.064 | -3.489 to -2.639 | 0.2032 | -0.2585 | -0.6122 to 0.09520 | **<0.0001** | 2.805 | 2.334 to 3.277 |
| **NEU (10³/µL)** | 0.1118 | 0.07939 to 0.1442 | 0.3010 | 0.0927 to 0.5093 | 0.1758 | 0.04625 to 0.3054 | **0.0038** | **0.0032** | -0.1892 | -0.3155 to -0.06291 | 0.3650 | -0.06400 | -0.1691 to 0.04108 | 0.0882 | 0.1252 | -0.01491 to 0.2653 |
| **NEU%** | 27.32 | 20.82 to 33.81 | 8.497 | 4.78 to 12.21 | 24.36 | 15.77 to 32.95 | **0.0088** | **0.0076** | 18.82 | 4.795 to 32.84 | >0.9999 | 2.955 | -8.713 to 14.62 | **0.0450** | -15.86 | -31.42 to -0.3057 |
| **LYM (10³/µL)** | 0.2507 | 0.1548 to 0.3466 | 3.030 | 2.298 to 3.762 | 0.4080 | 0.2619 to 0.5541 | **<0.0001** | **<0.0001** | -2.779 | -3.066 to -2.493 | 0.2864 | -0.1573 | -0.3954 to 0.08084 | **<0.0001** | 2.622 | 2.304 to 2.940 |
| **LYM%** | 56.19 | 47.03 to 65.35 | 86.67 | 81.13 to 92.20 | 61.12 | 44.78 to 77.46 | **0.0057** | **0.0048** | -30.48 | -51.85 to -9.108 | >0.9999 | -4.930 | -22.71 to 12.85 | **0.0328** | 25.55 | 1.840 to 49.25 |
| **MONO (10³/µL)** | 0.0358 | 0.02376 to 0.04784 | 0.052 | -0.0359 to 0.1399 | 0.0558 | 0.005798 to 0.1058 | 0.3861 | >0.9999 | -0.0162 | -0.06538 to 0.03298 | 0.6231 | -0.02 | -0.06538 to 0.03298 | >0.9999 | -0.0038 | -0.05836 to 0.05076 |
| **MONO%** | 8.718 | 5.723 to 11.71 | 1.461 | -0.7380 to 3.659 | 7.620 | 3.116 to 12.12 | **0.0325** | **0.0309** | 7.257 | 0.5914 to 13.92 | >0.9999 | 1.098 | -4.448 to 6.644 | 0.1212 | -6.159 | -13.55 to 1.236 |
| **EOS (10³/µL)** | 0.0038 | 0.002262 to 0.005338 | 0.023 | -0.0481 to 0.0941 | 0.008 | 0.0002957 to 0.0157 | 0.0573 | 0.0559 | -0.01920 | -0.03881 to 0.0004053 | >0.9999 | -0.0042 | -0.02051 to 0.01211 | 0.2488 | 0.015 | -0.00675 to 0.03675 |
| **EOS%** | 0.911 | 0.5863 to 1.236 | 0.7277 | -1.667 to 3.122 | 1.11 | 0.3099 to 1.91 | 0.6761 | >0.9999 | 0.1833 | -0.8780 to 1.245 | >0.9999 | -0.1990 | -1.082 to 0.6841 | >0.9999 | -0.3823 | -1.560 to 0.7951 |
| **BASO (10³/µL)** | 0.0306 | 0.01754 to 0.04366 | 0.0943 | 0.0123 to 0.1764 | 0.1136 | -0.08396 to 0.3112 | 0.1918 | 0.8053 | -0.06373 | -0.2131 to 0.08564 | 0.2766 | -0.083 | -0.2073 to 0.04129 | >0.9999 | -0.01927 | -0.1850 to 0.1465 |
| **BASO%** | 6.862 | 4.804 to 8.920 | 2.653 | 1,016 to 4,290 | 5.798 | 2.152 to 9.444 | 0.0931 | 0.0962 | 4.209 | -0.5899 to 9.007 | >0.9999 | 1.064 | -2.929 to 5.07 | 0.3972 | -3.145 | -8.468 to 2.179 |
| **RBC (10^6^/µL)** | 6.194 | 5.993 to 6.395 | 7.78 | 6,820 to 8,740 | 6.708 | 6.356 to 7.060 | **<0.0001** | **<0.0001** | -1.586 | -2.114 to -1.058 | **0.0198** | -0.5140 | -0.9533 to -0.07468 | **0.0005** | 1.072 | 0.4862 to 1.658 |
| **Hgb (g/dL)** | 9.625 | 9.323 to 9.927 | 11.3 | 10,80 to 11,80 | 10.25 | 9.528 to 10.98 | **0.0002** | **0.0001** | -1.675 | -2.475 to -0.8754 | 0.0670 | -0.6290 | -1.294 to 0.03628 | **0.0188** | 1.046 | 0.1590 to 1.933 |
| **HCT%** | 52.31 | 50.50 to 54.12 | 60.53 | 56.06 to 65.01 | 56.34 | 52.47 to 60.21 | **0.0006** | **0.0007** | -8.223 | -12.87 to -3.579 | **0.0396** | -4.030 | -7.894 to -0.1659 | 0.1336 | 4.193 | -0.9588 to 9.346 |
| **MCV (fL)** | 84.49 | 83.74 to 85.24 | 77.77 | 73.81 to 81.73 | 77.77 | 73.81 to 81.73 | **<0.0001** | <0.0001 | 6.723 | 4.582 to 8.864 | >0.9999 | 0.5300 | -1.251 to 2.311 | <0.0001 | -6.193 | -8.569 to -3.818 |
| **MCH (pg)** | 15.55 | 15.36 to 15.74 | 14.57 | 13.39 to 15.74 | 15.26 | 14.97 to 15.55 | **0.0006** | **0.0004** | 0.9833 | 0.4566 to 1.510 | 0.2848 | 0.2900 | -0.1483 to 0.7283 | **0.0180** | -0.6933 | -1.278 to -0.1089 |
| **MCHC (g/dL)** | 18.42 | 18.30 to 18.54 | 18.70 | 18.04 to 19.36 | 18.20 | 18.02 to 18.38 | **0.0056** | 0.0916 | -0.2800 | -0.5959 to 0.03588 | 0.1186 | 0.2200 | -0.04283 to 0.4828 | **0.0048** | 0.5000 | 0.1496 to 0.8504 |
| **RDW %CV** | 19.74 | 18.64 to 20.84 | 17.53 | 17.15 to 17.91 | 19.44 | 16.64 to 22.24 | 0.1620 | 0.4863 | 2.207 | -0.7422 to 5.156 | >0.9999 | 0.3000 | -2.154 to 2.754 | 0.4118 | -1.907 | -5.178 to 1.365 |
| **PLT (10³/µL)** | 488.4 | 440.1 to 536.7 | 670.3 | 548.7 to 791.9 | 487.6 | 359.3 to 615.9 | **0.0070** | **0.0079** | -181.7 | -318.1 to -45.72 | >0.9999 | 0.8000 | -112.5 to 114.1 | **0.0159** | 182.7 | 31.62 to 333.8 |
| **MPV (fL)** | 6.265 | 6.110 to 6.420 | 6.147 | 5.214 to 7.079 | 6.430 | 5.943 to 6.917 | 0.4116 | >0.9999 | 0.1183 | -0.4077 to 0.6444 | 0.9780 | -0.1650 | -0.6027 to 0.2727 | 0.6319 | -0.2833 | -0.8669 to 0.3003 |
| **PCT %** | 0.3061 | 0.2747 to 0.3375 | 0.4127 | 0.3048 to 0.5206 | 0.3114 | 0.2404 to 0.3824 | **0.0118** | **0.0120** | -0.1066 | -0.1912 to -0.02197 | >0.9999 | -0.0053 | -0.07569 to 0.06509 | **0.0325** | 0.1013 | 0.007414 to 0.1951 |
| **PDW 10(GSD)** | 17.30 | 16.97 to 17.63 | 17.43 | 16.67 to 18.19 | 17.62 | 16.79 to 18.45 | 0.5342 | >0.9999 | -0.1333 | -1.042 to 0.7749 | 0.8156 | -0.3200 | -1.076 to 0.4357 | >0.9999 | -0.1867 | -1.194 to 0.8209 |

**Supplementary table 2 S2: blood cell counts of control and weekly µCT-scanned healthy and animals with bleomycin-induced lung inflammation and fibrosis.**

WBC=white blood cells; NEU=neutrophils; LYM=lymphocytes; MONO=monocytes; EOS=eosinophils; BASO=basophils; RBC=red blood cells; Hgb=hemoglobin; HCT=hematocrit; MCV=mean corpuscular (cell) volume; MCH=mean corpuscular hemoglobin; MCHC=mean corpuscular hemoglobin concentration; RDW=red cell distribution width; PLT=platelets; MPV=mean platelet volume; PCT=plateletcrit

|  | **Exp3 - healthy** | | | | | | |
| --- | --- | --- | --- | --- | --- | --- | --- |
|  | **Healthy control** | | **Healthy weekly CT** | |  | | |
|  | **mean** | **95% CI** | **mean** | **95% CI** | **p-value** | **mean difference** | **95% CI** |
| **WBC (10³/µL)** | 0.6140 | 0.04885 to 1.179 | 0.2630 | 0.07476 to 0.4512 | **0.0431** | **-0.3510** | -0.6859 to -0.01608 |
| **NEU (10³/µL)** | 0.1800 | 0.04855 to 0.3114 | 0.1115 | 0.05936 to 0.1636 | 0.0858 | -0.06850 | -0.1510 to 0.01396 |
| **NEU%** | 29.93 | 21.78 to 38.08 | 44.30 | 30.89 to 57.71 | **0.0405** | **14.37** | 0.9147 to 27.82 |
| **LYM (10³/µL)** | 0.4057 | 0.03771 to 0.7736 | 0.1208 | -0.02367 to 0.2652 | **0.0244** | **-0.2849** | -0.5149 to -0.05496 |
| **LYM%** | 66.00 | 58.72 to 73.28 | 42.75 | 21.58 to 63.92 | **0.0335** | **-23.25** | -43.81 to -2.695 |
| **MONO (10³/µL)** | 0.01733 | -0.02720 to 0.06186 | 0.01550 | 0.008443 to 0.02256 | 0.8474 | -0.001833 | -0.02509 to 0.02142 |
| **MONO%** | 2.450 | -1.817 to 6.717 | 6.635 | 2.084 to 11.19 | 0.0770 | 4.185 | -0.6591 to 9.029 |
| **EOS (10³/µL)** | 0.001667 | -0.002128 to 0.005461 | 0.007000 | -0.007983 to 0.02198 | 0.3862 | 0.005333 | -0.009112 to 0.01978 |
| **EOS%** | 0.3303 | -0.2564 to 0.9170 | 3.114 | -3.988 to 10.22 | 0.3404 | 2.784 | -4.010 to 9.578 |
| **BASO (10³/µL)** | 0.009667 | -0.02119 to 0.04053 | 0.008250 | 0.001836 to 0.01466 | 0.835 | -0.001417 | -0.01802 to 0.01518 |
| **BASO%** | 1.264 | -1.915 to 4.444 | 3.190 | 2.242 to 4.138 | **0.0425** | **1.926** | 0.09624 to 3.755 |
| **RBC (10^6^/µL)** | 8.267 | 7.954 to 8.579 | 6.435 | 4.554 to 8.316 | **0.0477** | **-1.832** | -3.637 to -0.02679 |
| **Hgb (g/dL)** | 11.07 | 10.44 to 11.69 | 8.678 | 6.238 to 11.12 | **0.0476** | **-2.389** | -4.742 to -0.03664 |
| **HCT%** | 61.03 | 58.26 to 63.80 | 48.80 | 34.71 to 62.89 | 0.0678 | -12.23 | -25.77 to 1.305 |
| **MCV (fL)** | 73.87 | 71.99 to 75.75 | 75.90 | 74.91 to 76.89 | **0.0112** | **2.033** | 0.6998 to 3.367 |
| **MCH (pg)** | 13.40 | 12.65 to 14.15 | 13.50 | 13.16 to 13.84 | 0.6268 | 0.1000 | -03967 to 0.5967 |
| **MCHC (g/dL)** | 18.17 | 17.65 to 18.68 | 17.78 | 17.23 to 18.32 | 0.1423 | -0.3917 | -0.9702 to 0.1869 |
| **RDW %CV** | 17.83 | 15.10 to 20.57 | 19.33 | 17.62 to 21.03 | 0.1309 | 1.492 | -0.6327 to 3.616 |
| **PLT (10³/µL)** | 735.7 | 534.4 to 936.9 | 347.8 | 255.7 to 439.8 | **0.0007** | **-387.9** | -521.6 to -254.3 |
| **MPV (fL)** | 6.443 | 6.083 to 6.804 | 8.005 | 7.459 to 8.551 | **0.0008** | **1.562** | 1.010 to 2.114 |
| **PCT %** | 0.4743 | 0.3252 to 0.6235 | 0.2770 | 0.2167 to 0.3373 | **0.0030** | **-0.1973** | -0.2916 to -0.1031 |
| **PDW 10(GSD)** | 17.30 | 16.64 to 17.96 | 19.28 | 17.47 to 21.08 | **0.0340** | **1.975** | 0.2217 to 3.728 |

**Supplementary table 3 S3: blood cell counts of control and weekly µCT-scanned healthy animals.**

WBC=white blood cells; NEU=neutrophils; LYM=lymphocytes; MONO=monocytes; EOS=eosinophils; BASO=basophils; RBC=red blood cells; Hgb=hemoglobin; HCT=hematocrit; MCV=mean corpuscular (cell) volume; MCH=mean corpuscular hemoglobin; MCHC=mean corpuscular hemoglobin concentration; RDW=red cell distribution width; PLT=platelets; MPV=mean platelet volume; PCT=plateletcrit

|  | **Exp4 - healthy** | | | | | | |
| --- | --- | --- | --- | --- | --- | --- | --- |
|  | **Healthy control** | | **Healthy one CT** | |  | | |
|  | **mean** | **95% CI** | **mean** | **95% CI** | **p-value** | **mean difference** | **95% CI** |
| **WBC (10³/µL)** | 0.5217 | 0.3006 to 0.7427 | 0.5760 | 0.2932 to 0.8588 | 0.6908 | -0.05433 | -0.3535 to 0.2449 |
| **NEU (10³/µL)** | 0.08833 | 0.04940 to 0.1273 | 0.1020 | 0.05689 to 0.1471 | 0.5545 | -0.01367 | -0.06402 to 0.03669 |
| **NEU%** | 17.18 | 13.13 to 21.24 | 18.46 | 11.78 to 25.14 | 0.6575 | -1.277 | -7.576 to 5.023 |
| **LYM (10³/µL)** | 0.4050 | 0.2179 to 0.5921 | 0.4400 | 0.1711 to 0.789 | 0.775 | -0.03500 | -0.3038 to 0.2338 |
| **LYM%** | 77.35 | 72.84 to 81.86 | 74.7 | 65.54 to 83.86 | 0.4748 | 2.650 | -5.388 to 10.69 |
| **MONO (10³/µL)** | 0.008333 | 0.004049 to 0.01262 | 0.01800 | 0.007611 to 0.02839 | **0.0332** | **-0.009667** | -0.01837 to -0.0009632 |
| **MONO%** | 1.633 | 0.9012 to 2.365 | 3.360 | 0.1238 to 6.596 | 0.1504 | -1.727 | -4.211 to 0.7577 |
| **EOS (10³/µL)** | 0.01167 | -0.0006017 to 0.02394 | 0.00800 | -0.002389 to 0.01839 | 0.5727 | 0.003667 | -0.01051 to 0.01784 |
| **EOS%** | 1.933 | 0.5688 to 3.298 | 1.800 | -0.3524 to 3.952 | 0.8871 | 0.1333 | -1.933 to 2.199 |
| **BASO (10³/µL)** | 0.003333 | -0.002086 to -0.007106 | 0.00400 | -0.007106 to 0.01511 | 0.8801 | -0.0006667 | -0.01039 to 0.009055 |
| **BASO%** | 0.6500 | -0.2122 to 1.512 | 1.060 | -0.5072 to 2.627 | 0.5315 | -0.4100 | -1.836 to 1.016 |
| **RBC (10^6^/µL)** | 5.390 | 4.232 to 6.548 | 5.590 | 4.379 to 6.801 | 0.7599 | -0.2000 | -1.636 to 1.236 |
| **Hgb (g/dL)** | 7.883 | 6.184 to 9.583 | 8.240 | 6.424 to 10.06 | 0.7130 | -0.3567 | -2.482 to 1.768 |
| **HCT%** | 25.25 | 18.89 to 31.61 | 27.04 | 20.48 to 33.60 | 0.6182 | -1.790 | -9.635 to 6.055 |
| **MCV (fL)** | 46.60 | 44.73 to 48.47 | 48.22 | 46.73 to 49.71 | 0.1189 | -1.620 | -3.746 to 0.5064 |
| **MCH (pg)** | 14.60 | 14.42 to 14.78 | 14.78 | 14.62 to 14.94 | 0.0823 | -0.1800 | 0.3882 to 0.02824 |
| **MCHC (g/dL)** | 31.38 | 30.16 to 32.60 | 30.66 | 29.82 to 31.50 | 0.2528 | 0.7233 | -0.6157 to 0.5919 |
| **RDW %CV** | 13.20 | 12.59 to 13.81 | 14.12 | 13.58 to 14.66 | **0.0169** | **-0.9200** | -1.631 to -0.2085 |
| **PLT (10³/µL)** | 531.0 | 369.9 to 665.1 | 627.2 | 479.8 to 774.6 | 0.2315 | -96.20 | -265.8 to 73.41 |
| **MPV (fL)** | 7.283 | 6.160 to 8.407 | 7.060 | 6.026 to 8.094 | 0.7131 | 0.2233 | -1.108 to 1.555 |
| **PCT %** | 0.3750 | 0.3139 to 0.4361 | 0.4440 | 0.3139 to 0.5741 | 0.1992 | -0.06900 | -0.1817 to 0.04365 |

**Supplementary table 4 S4: blood cell counts of control and healthy animals after a single µCT scan.**

WBC=white blood cells; NEU=neutrophils; LYM=lymphocytes; MONO=monocytes; EOS=eosinophils; BASO=basophils; RBC=red blood cells; Hgb=hemoglobin; HCT=hematocrit; MCV=mean corpuscular (cell) volume; MCH=mean corpuscular hemoglobin; MCHC=mean corpuscular hemoglobin concentration; RDW=red cell distribution width; PLT=platelets; MPV=mean platelet volume; PCT=plateletcrit

|  | **Exp5 - healthy** | | | | | | | | | | | | | | | |
| --- | --- | --- | --- | --- | --- | --- | --- | --- | --- | --- | --- | --- | --- | --- | --- | --- |
|  | **Healthy weekly sedation** | | **Healthy control** | | **Healthy weekly CT** | |  | **Healthy WS vs healthy CO** | | | **Healthy WS vs healthy weekly CT** | | | **Healthy CO vs healthy weekly CT** | | |
|  | **mean** | **95% CI** | **mean** | **95% CI** | **mean** | **95% CI** | **p-value** | **p-adj value** | **mean difference** | **95% CI** | **p-adj value** | **mean difference** | **95% CI** | **p-adj value** | **mean difference** | **95% CI** |
| **WBC (10³/µL)** | 1.650 | 1.168 to 2.132 | 1.178 | 0.7948 to 1.1560 | 1.178 | 0.7774 to 1.578 | 0.0921 | 0.1642 | -0.4725 | -1.085 to 0.1401 | 0.1969 | -0.4722 | -1.115 to 0.1704 | >0.9999 | 0.003000 | -0.6423 to 0.6429 |
| **NEU (10³/µL)** | 0.3870 | 0.2761 to 0.4979 | 0.3030 | 0.2480 to 0.3580 | 0.2988 | 0.2259 to 0.3717 | 0.1260 | 0.2407 | 0.3870 | -0.2051 to 0.03706 | 0.2398 | -0.08820 | -0.2152 to 0.03877 | >0.9999 | -0.004200 | -0.1312 to 0.1228 |
| **NEU%** | 24.03 | 17.46 to 30.60 | 27.42 | 19.84 to 35 | 26.68 | 17.23 to 36.13 | 0.6895 | >0.9999 | 3.383 | -7.622 to 14.39 | >0.9999 | 2.647 | -8.896 to 14.19 | >0.9999 | -0.7367 | -12.28 to 10.81 |
| **LYM (10³/µL)** | 1.127 | 0.7268 to 1.527 | 0.8027 | 0.4856 to 1.120 | 0.7896 | 0.3975 to 1.182 | 0.1900 | 0.3511 | -0.3242 | -0.8516 to 0.2033 | 0.3593 | -0.3372 | -0.8904 to 0.2159 | >0.9999 | -0.0131 | -0.5662 to 0.5401 |
| **LYM%** | 67.40 | 58.08 to 76.72 | 66.42 | 58.63 to 74.20 | 65.46 | 52.40 to 78.52 | 0.9375 | >0.9999 | -0.9833 | -14.97 to 13.00 | >0.9999 | -1.940 | -16.61 to 21.73 | >0.9999 | -0.9567 | -15.62 to 13.71 |
| **MONO (10³/µL)** | 0.05650 | 0.02252 to 0.09048 | 0.03383 | 0.02111 to 0.04656 | 0.05200 | 0.005235 to 0.09876 | 0.3842 | 0.5852 | -0.02267 | -0.06793 to 0.02260 | >0.9999 | -0.004500 | -0.05198 to 0.04298 | 0.9480 | 0.01817 | -0.02931 to 0.06564 |
| **MONO%** | 3.641 | 1.177 to 6.105 | 2.955 | 2.204 to 3.706 | 4.710 | 0.4641 to 8.956 | 0.4820 | >0.9999 | -0.6858 | -4.363 to 2.992 | >0.9999 | 1.069 | -2.788 to 4.926 | 0.7096 | 1.755 | -2.102 to 5.612 |
| **EOS (10³/µL)** | 0.02567 | 0.01723 to 0.03411 | 0.02117 | 0.009511 to 0.03282 | 0.01800 | 0.006860 to 0.02914 | 0.4246 | >0.9999 | -0.004500 | -0.01940 to 0.01040 | 0.6110 | -0.007667 | -0.02329 to 0.007959 | >0.9999 | -0.003167 | -0.01879 to 0.01246 |
| **EOS%** | 1.575 | 1.119 to 2.031 | 1.770 | 1.019 to 2.521 | 1.493 | 0.9352 to 2.050 | 0.6965 | >0.9999 | 0.3204 | -0.6755 to 1.066 | >0.9999 | -0.08220 | -0.9955 to 0.8311 | >0.9999 | -0.2775 | -1.191 to 0.6358 |
| **BASO (10³/µL)** | 0.05383 | 0.03005 to 0.07762 | 0.02950 | -0.001876 to 0.06088 | 0.01880 | 0.006004 to 0.03160 | 0.0627 | 0.2680 | -0.02433 | -0.06056 to 0.01189 | 0.0755 | 0.05383 | -0.07303 to 0.002963 | >0.9999 | -0.01070 | -0.04870 to 0.02730 |
| **BASO%** | 3.317 | 1.855 to 4.779 | 1.428 | 1.010 to 1.847 | 1.658 | 0.4805 to 2.835 | **0.0117** | **0.0171** | **-1.888** | -3.463 to -0.3139 | **0.0488** | **-1.659** | -3.310 to -0.007812 | >0.9999 | 0.2293 | -1.422 to 1.881 |
| **RBC (10^6^/µL)** | 8.405 | 8.082 to 8.728 | 8.005 | 7.596 to 8.414 | 8.304 | 7.871 to 8.737 | 0.1603 | 0.2046 | 0.2990 | -0.9502 to 0.1502 | >0.9999 | -0.1010 | -0.6780 to 0.4760 | 0.5426 | 0.2990 | -0.2780 to 0.8760 |
| **Hgb (g/dL)** | 11.15 | 10.66 to 11.64 | 10.68 | 10.10 to 11.27 | 10.80 | 9.588 to 12.00 | 0.5739 | 0.9591 | -0.4667 | -1.688 to 0.7547 | >0.9999 | -0.3533 | -1.575 to 0.8681 | >0.9999 | 0.1133 | -1.108 to 1.335 |
| **HCT%** | 60.87 | 58.64 to 63.09 | 58.28 | 55.12 to 61.45 | 60.68 | 57.76 to 63.60 | 0.1877 | 0.2989 | -2.583 | -6.565 to 1.398 | >0.9999 | -0.1867 | -4.362 to 3.989 | 0.4233 | 2.3970 | -1.779 to 6.572 |
| **MCV (fL)** | 72.42 | 71.77 to 73.06 | 72.80 | 72.02 to 73.58 | 73.12 | 72.37 to 73.87 | 0.2431 | 0.9942 | 0.3833 | -0.6521 to 1.419 | 0.3006 | 0.7033 | -0.7660 to 1.406 | >0.9999 | 0.3200 | -0.7660 to 1.406 |
| **MCH (pg)** | 13.27 | 13.06 to 13.47 | 13.35 | 13.21 to 13.49 | 13.54 | 13.35 to 13.73 | **0.0459** | >0.9999 | 0.08333 | -0.1753 to 0.3420 | **0.0480** | **0.2733** | 0.002087 to 0.5446 | 0.2331 | 0.1900 | -0.08125 to 0.4612 |
| **MCHC (g/dL)** | 18.35 | 18.19 to 18.51 | 18.35 | 18.26 to 18.44 | 18.50 | 18.38 to 18.62 | 0.0868 | >0.9999 | 0.000 | -0.1828 to 0.1828 | 0.1552 | 0.1500 | -0.4172 to 0.3417 | 0.1552 | 0.1500 | -0.4172 to 0.3417 |
| **RDW %CV** | 18.25 | 17.41 to 19.09 | 17.38 | 15.82 to 18.94 | 18.12 | 17.96 to 18.28 | 0.3137 | 0.4794 | -0.8667 | -2.453 to 0.7198 | >0.9999 | -0.1300 | -1.794 to 1.534 | 0.7465 | 0.7367 | -0.9272 to 2.401 |
| **PLT (10³/µL)** | 710.0 | 643.0 to 777.0 | 696.7 | 638.3 to 755.1 | 650.2 | 597.8 to 702.6 | 0.2168 | >0.9999 | -13.33 | -100.3 to 73.63 | 0.2894 | -59.80 | -151.0 to 31.41 | 0.5636 | -46.47 | -137.7 to 44.74 |
| **MPV (fL)** | 5.972 | 5.670 to 6.274 | 6.528 | 5.839 to 7.218 | 6.520 | 6.063 to 6.977 | 0.1043 | 0.1805 | 0.5567 | -0.1833 to 1.297 | 0.2263 | 0.5483 | -0.2277 to 1.324 | >0.9999 | -0.008333 | -0.7844 to 0.7677 |
| **PCT %** | 0.4245 | 0.3748 to 0.4742 | 0.4520 | 0.4271 to 0.4769 | 0.4290 | 0.3500 to 0.5080 | 0.4844 | 0.7847 | 0.02750 | -0.03684 to 0.09184 | >0.9999 | 0.004500 | -0.06743 to 0.07643 | >0.9999 | -0.02300 | -0.09493 to 0.04893 |
| **PDW 10(GSD)** | 17.87 | 17.46 to 18.27 | 18.73 | 16.57 to 20.89 | 18.38 | 16.71 to 20.04 | 0.5710 | 0.9021 | 0.8667 | -1.341 to 3.075 | >0.9999 | 0.5083 | -1.960 to 2.977 | >0.9999 | -0.3583 | -2.827 to 2.110 |

**Supplementary table 5 S5: blood cell counts of control and weekly µCT-scanned healthy animals.**

WBC=white blood cells; NEU=neutrophils; LYM=lymphocytes; MONO=monocytes; EOS=eosinophils; BASO=basophils; RBC=red blood cells; Hgb=hemoglobin; HCT=hematocrit; MCV=mean corpuscular (cell) volume; MCH=mean corpuscular hemoglobin; MCHC=mean corpuscular hemoglobin concentration; RDW=red cell distribution width; PLT=platelets; MPV=mean platelet volume; PCT=plateletcrit
